# Functional Cliques in Developmentally Correlated Neural Networks

**DOI:** 10.1101/064147

**Authors:** S. Luccioli, A. Barzilai, E. Ben-Jacob, P. Bonifazi, A. Torcini

**Affiliations:** CNR - Consiglio Nazionale delle Ricerche - Istituto dei Sistemi Complessi, 50019 Sesto Fiorentino, Italy; INFN - Istituto Nazionale di Fisica Nucleare - Sezione di Firenze, 50019 Sesto Fiorentino, Italy: Joint Italian-Israeli Laboratory on Integrative Network Neuroscience, Tel Aviv University, Ramat Aviv, Israel.; Joint Italian-Israeli Laboratory on Integrative Network Neuroscience, Tel Aviv University, Ramat Aviv, Israel; Department of Neurobiology, George S. Wise Faculty of Life Sciences and Sagol School of Neuroscience, Tel Aviv University, Ramat Aviv, Israel.; Joint Italian-Israeli Laboratory on Integrative Network Neuroscience, Tel Aviv University, Ramat Aviv, Israel: Beverly and Sackler Faculty of Exact Sciences School of Physics and Astronomy, Tel Aviv University, Ramat Aviv, Israel.; Joint Italian-Israeli Laboratory on Integrative Network Neuroscience, Tel Aviv University, Ramat Aviv, Israel; Beverly and Sackler Faculty of Exact Sciences School of Physics and Astronomy, Tel Aviv University, Ramat Aviv, Israel; Computational Neuroimaging Lab, BioCruces Health Research Institute, Hospital Universitario Cruces, Plaza de Cruces, s/n E-48903, Barakaldo, Spain.; Aix Marseille Univ, Inserm, INMED, Institute de Neurobiologie de la Méditerranée and INS, Institut de Neurosciences des Systémes, Marseille, France; Aix-Marseille Université, Université de Toulon, CNRS, CPT, UMR 7332, 13288 Marseille, France; CNR - Consiglio Nazionale delle Ricerche - Istituto dei Sistemi Complessi, 50019 Sesto Fiorentino, Italy: Joint Italian-Israeli Laboratory on Integrative Network Neuroscience, Tel Aviv University, Ramat Aviv, Israel.

**Author notes:** These authors are joint senior authors on this work (P. Bonifazi, A. Torcini).

## Abstract

We consider a sparse random network of excitatory leaky integrate-andfire neurons with short-term synaptic depression. Furthermore to mimic the dynamics of a brain circuit in its first stages of development we introduce for each neuron correlations among in-degree and out-degree as well as among excitability and the corresponding total degree, We analyze the influence of single neuron stimulation and deletion on the collective dynamics of the network. We show the existence of a small group of neurons capable of controlling and even silencing the bursting activity of the network. These neurons form a *functional clique* since only their activation in a precise order and within specific time windows is capable to ignite population bursts.

## 1.1 Introduction

The relationship among brain functional and structural connectivity and neuronal activity is one of the subject of major interest in neuroscience [1, 2, 3, 4, 5]. Furthermore, this research line is also strongly related to nonlinear dynamics themes regarding the influence of topology on the emergence of specific microscopic and collective dynamics [6, 7, 8].

On one side, it is nowdays clear that a specific topology is not sufficient to guarantee an unequivocal dynamical behaviour in the neural network [9]. On the other side, in the last two years experimental and numerical evidences have been indicating that neuronal ensembles or cell assemblies (*cliques*) are the emergent functional units of cortical activity [10, 11, 12, 13], shaping spontaneous and stimuli/task-evoked responses. Interstingly, several experimental studies have revealed that single neurons can have a relevant role in shaping neuronal dynamics in brain circuits [14, 15, 16, 17, 18, 19, 20, 21]. Therefore it is of major importance to understand how do neuronal cliques emerge and how they do operate in relation to single neuron dynamics. In this framework we could link the single neuron firing to the emergence of neuronal cliques, in a sort of hierarchical modular approach where high order dynamics (i.e involving large number of neurons) can be strongly impacted by single neuron manipulations.

Following this line of experimental and theoretical evidence, and in particular the analysis performed in [21], we have derived a numerical model displaying collective oscillations, similar to the ones observed in the hyppocampus and in the neocortex at the early stage of development [22]. In particular, we have analyzed the influence of single neuron stimulation/deletion on the dynamics of such a network in presence of developmentally inspired constraints. Our analysis reveals that a small group of critical neurons, organized in a *clique*, is capable to control the bursting behaviour of the whole network. These neurons are *hubs* in a *functional* sense, because their relevance is not related to high intrinsic connectivity, but to their precise sequential and coordinated activation before a bursting event. As a consequence, if a perturbation is applied to any critical neuron of the *functional clique*, through their stimulation or the deletion, their sequential activation can be interrupted, thus leading to dramatic consequence at a network level, with the arrest of the collective oscillations.

The studied model and the methods employed to analyze the data are introduced in Sect. 1.2, with particular emphasis on functional connectivity. Sect. 1.3 reports the results of the numerical experiments performed at the single neuron level in our network and of the functional analysis. Finally a brief discussion is reported in Sect. 1.4.

## 1.2 Model and Methods

We consider a network of *N* excitatory Leaky Integrate-and-fire (LIF) neurons, interacting via synaptic currents regulated by short-term-plasticity according to the model introduced in [23]. For this model, the evolution of the membrane potential *V_i_* of the neuron *i* is given by

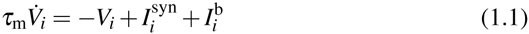

where *τ*_m_ is the membrane time constant, 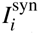 is the synaptic current received by neuron *i* from all its presynaptic inputs and 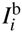 represents its level of intrinsic excitability, encompassing single neuron properties as well as excitatory inputs arriving from distal areas. The currents are measured in voltage units (mV), since the membrane input resistance is incorporated into the currents.

Whenever the membrane potential *V_i_*(*t*) reaches the threshold value *V*_th_, it is reset to *V*_r_, and a *δ*-spike is sent towards the postsynaptic neurons. Accordingly, the spike-train *S_j_*(*t*) produced by neuron *j*, is defined as,

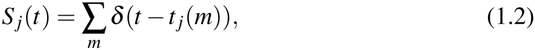

where *t_j_*(*m*) represent the *m*-th spike time emission of neuron *j*. The transmission of the spike train *S_j_* to the efferent neurons is mediated by the synaptic evolution. In particular, by following [24] the state of the synapse between the *j*th presynaptic neuron and the *i*th postsynaptic neuron is described by three adimensional variables, *X_ij_*, *Y_ij_*, and *Z_ij_*, representing the fractions of synaptic resources in the recovered, active, and inactive state, respectively.

Since these three variables should satisfy the constraint *X_ij_* + *Y_ij_* + *Z_ij_* = 1, it is sufficient to provide the time evolution for two of them, namely

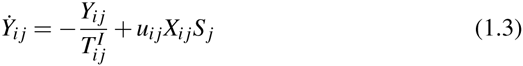

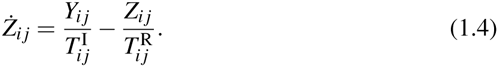

Only the active transmitters react to the incoming spikes *S_j_* and the adimensional parameters *u_ij_* tunes their effectiveness. The decay times of the postsynaptic current are given by 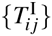, while 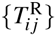 represent the recovery times from the synaptic depression.

The synaptic current is expressed as the sum of all the active transmitters (post-synaptic currents) delivered to neuron *i*

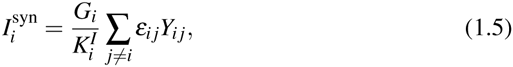

where *G_i_* is the coupling strength, while ε_*ij*_ is the connectivity matrix whose entries are set equal to 1 (0) if the presynaptic neuron *j* is connected to (disconnected from) the postsynaptic neuron *i*. At variance with [23], we assume that the coupling strengths are the same for all the synapses afferent to a certain neuron *i*.

In particular, we study the case of excitatory coupling between neurons, i.e. *G_i_* > 0. Moreover, we consider a sparse network made of *N* = 200 neurons where the *i*-th neuron has 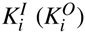 afferent (efferent) synaptic connections distributed as in a directed Erdös-Rényi graph with average in-degree 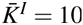, as a matter of fact also the average out-degree was 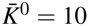. The sum appearing in (1.5) is normalized by the input degree 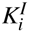 to ensure homeostatic synaptic inputs [25, 26].

The intrinsic excitabilities of the single neurons 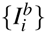 are randomly chosen from a flat distribution of width 0.45 mV centered around the value *V*_th_ = 15 mV, by further imposing that 5% of neurons are above threshold in order to observe a bursting behaviour in the network. For the other parameters, in analogy with Ref. [23], we use the following set of values: *τ*_m_ = 30 ms,*V*_r_ = 13.5 mV, *V*_th_ = 15 mV. The synaptic parameters 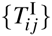, 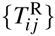, {*u_ij_*} and {*G_i_*} are Gaussian distributed with averages 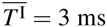, 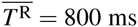, 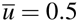 and 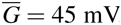, respectively, and with standard deviation equal to the half of the average.

### 1.2.1 Correlations

In this paper, we intend to mimic neural networks at the first stage of development. In such networks mature and young neurons are present at the same time, and this is reflected in the variability of the structural connectivities and of the intrinsic excitabilities. In particular, experimental observations indicate that younger cells have a more pronounced excitability [27, 28], while mature cells exhibit a higher number of synaptic inputs [21, 29]. Thus suggesting that the number of afferent and efferent synaptic connections [21, 29, 30] as well as their level of hyperpolarization [31] are positively correlated with the maturation stage of the cells. Therefore, we consider a network including the following two types of correlations:

- setup T1: a positive correlation between the in-degree 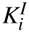 and out-degree 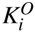 of each neuron;
- setup T2: a negative correlation between the intrinsic neuronal excitability and the total connectivity 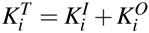 (in-degree plus out-degree) ;

Correlation of type T1 is obtained by generating randomly two pools of *N* input and output degrees from an Erdös-Rényi distribution with average degree equal to 10. The degrees are ordered within each pool and then assigned to *N* neurons in order to obtain a positive correlation between 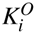 and 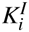.

Correlation of type T2 imposes a negative correlation between excitability 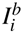 and the total degree of the single neuron 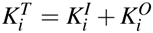. To generate this kind of correlation the intrinsic excitabilities are randomly generated, as explained above, and then assigned to the various neurons accordingly to their total connectivities 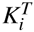, thus to ensure an inverse correlation between 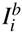 and 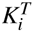.

### 1.2.2 Functional Connectivity

In order to highlight statistically significant time-lagged activations of neurons, for every possible neuronal pair, we measure the cross-correlation between their spike time series. On the basis of this cross-correlation we eventually assign a directed functional connection among the two considered neurons, similarly to what reported in [21, 32] for calcium imaging studies.

For every neuron, the action potentials timestamps were first converted into a binary time series with one millisecond time resolution, where ones (zeros) marked the occurrence (absence) of the action potentials. Given the binary time series of two neurons *a* and *b*, the cross correlation was then calculated as follows:

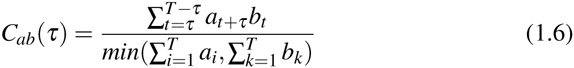

where {*a_t_}*,{*b_t_*} represented the considered time series and *T* was their total duration. Whenever *C_ab_*(*τ*) presented a maximum at some finite time value *τ_max_* a functional connection was assigned between the two neurons: for *τ_max_* < 0 (*τ_max_* > 0) directed from *a* to *b* (from *b* to *a*). A directed functional connection cannot be defined for an uniform cross-correlation corresponding to uncorrelated neurons or for synchronous firing of the two neurons associated to a Gaussian *C_ab_*(*τ*) centered at zero. To exclude the possibility that the cross correlation could be described by a Gaussian with zero mean or by a uniform distribution we employed both the Student’s t-test and the Kolmogorov-Smirnov test with a level of confidence of 5%. The functional out-degree 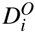 (in-degree 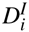) of a neuron *i* corresponded to the number of neurons which were reliably activated after (before) its firing.

For more deiails on the model and methods employed to perform the reported analysis see [13].

## 1.3 Results

In this paper we intend to mimic an immature neuronal network at a certain stage of its initial development, in analogy with the one examined in the experimental work on developmental hippocampal circuits [21]. As discussed in [21, 22], at early post-natal stages such networks are characterized by the excitatory action of GABAergic transmission and the presence of synchronized network events, as largely documented in central and peripheral nervous circuits.

Therefore, we consider a network model composed of only excitatory neurons and displaying bursting activity. A minimal model to mimic the experimentally described stereotypical/characteristic condition of developing neuronal networks [33] is the one introduced by Tsodyks-Uziel-Markram (TUM) [23] with purely excitatory synapses. Since this model (described in details in Sect. 1.2) is known to display an alternance of short periods of quasi-synchronous firing (*population bursts*, PBs) and long time intervals of asynchronous firing. Furthermore, we consider a network with embedded correlations of type T1 and T2, this in order to account for the presence at the same time of younger and older neurons. This presence can be modeled by considering correlations among the in-degree and out-degree of each cell (type T1) as well as among their intrinsic excitability and connectivity (type T2), as already explained in the previous Section. The network activity was characterized by bursts of duration ≃ 24 ms and with interburst intervals ≃ 500 ms.

### 1.3.1 Single neuron stimulation and deletion experiments

In the developing hippocampus it has been shown that the stimulation of specific single neurons can drastically reduce the frequency of the PBs [3, 21], or even silence the collective activity. More specifically, the stimulation consisted of current injection capable of inducing sustained high firing regime of the stimulated neuron over a period of a few minutes.

Inspired by this experimental protocol, we test the impact of *single neuron stimulation* (SNS) on the occurrence of PBs in our network model. SNS was achieved by adding abruptly a DC current term to the considered neuron. We report in Fig. 1.1 A-B the stimulation protocol for a specific neuron capable of suppressing the occurrence of PBs for all the duration of the SNS (in this case limited to 4.2 s). The SNS process is totally reversible, i.e. when the stimulation is interrupted the firing rate of the cell and the PBs frequency returns to the pre-stimulation control level.

**Fig. 1.1.**
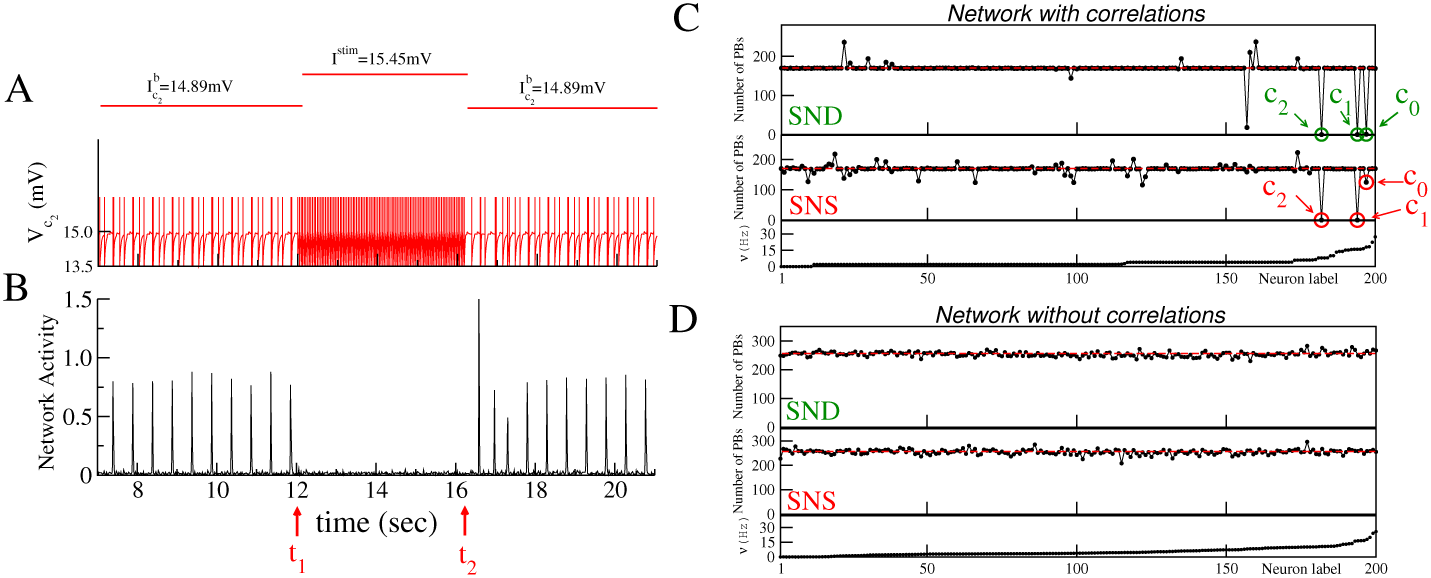
(A), (B) Sketch of a SNS experiment for a network with type T1 plus T2 correlations. The neuron *c*_2_ is stimulated with a DC step for a time interval *Δt* = *t*_2_ − *t*_1_ (see the red line on the top panel). In (A) the membrane potential *V_c_2__* of the stimulated neuron is shown (firing times are signaled by vertical bars of height 1.5 mV), while in (B) the network activity is reported. (C) and (D) refer to correlated and uncorrelated networks, respectively, and display the number of PBs emitted in a time window *Δt* = 84 s during SND and SNS experiments with *I*^stim^ = 15.45 mV. The horizontal dashed lines refer to the average number of PBs emitted in a time interval *Dt* = 84 s during a control experiment when no stimulation is applied (the amplitude of the fluctuations is smaller than the symbols). Neurons are ordered accordingly to their average firing rates *n* measured in control condition (plotted in the bottom panels). The critical neurons *c*_0_, *c*_1_, *c*_2_, signaled by green (red) circles, are able to strongly affect the bursting during SND (SNS). During SNS experiments each neuron *i* was stimulated with a DC step switching its excitability from 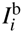 to *I*^stim^ for an interval *Δt* = 84 s. The data refer to *I*^stim^ = 15.45 mV and *N* = 200 neurons.

In order to measure the impact of SNS on the network dynamics, we consider the relative variation of the PB frequency with respect to the control conditions (i.e. in absence of any stimulation). In Figs. 1.1 C-D the impact of a SNS on the PBs frequency is reported for a classical Erdös-Rényi network (no correlations) and a network with embedded correlations T1 plus T2. It is clear that SNS has much more impact on the correlated network, than in the uncorrelated one. In particular for two neurons, indicated as neuron *c*_1_ and *c*_2_, SNS performed with stimulation current *I*^stim^ = 15.45 mV, is able to suppress the occurrence of PBs during the stimulation period. Furthermore in other 7 cases SNS reduces the collective activity ≃ 30%, in the specific case of neuron *c*_0_ (whose relevance is discussed in the following) the reduction was of ≃ 27%.

On the contrary, in the network without correlations, the SNS had only marginal effect on the population activity, although the distributions of the firing rates in the correlated and uncorrelated network are extremely similar (under control conditions) as shown in the bottom panels of Fig. 1.1 C and D.

In [23] it has been shown that the elimination of a pool of neurons from an uncorrelated TUM network induced a strong reduction of the population bursts. In our analysis we repeat such numerical experiment with single cell resolution, i.e. we consider the influence of *single neuron deletion*, SND, on the network response. The results reported in Fig. 1.1 C-D, clearly show almost no effect for the uncorrelated network. However, SND blocks the deliver of PBs in the network with correlations *T*1 plus *T*2, in three cases. It should be noticed that two over three neurons were critical also fro the SNS, while the third critical neuron has a higher frequency and it is denoted as *c*_0_.

The most critical neurons are all unable to fire if isolated from the network, i.e. they are all below threshold, apart neuron *c*_0_, which has 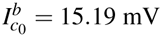. Therefore we observe that SND or SNS are capable to silence the network and that this occurs only for a quite limited number of neurons, which have reasonably high *I^b^* but low *K^T^* (*c*_0_, with *K^T^* =6, is ranked as the second less connected neuron of the network).

### 1.3.2 The clique of functional hubs

The detailed investigation of the burst events revealed that each PB is always preceeded by the firing of the 3 critical neurons in the following order: *c*_0_ → *c*_1_ → *c*_2_ → PB, as it is shown in Fig. 1.2. The neuron *c*_0_, which is the only one supra-threshold, fires first followed by the others (sub-threshold) and this trigger the onset of the PB. Furthermore, the functional connectivity analysis of the neurons in the network confirms that *c*_0_, *c*_1_, and *c*_2_ have quite high functional output degree *D^O^* ≃ 170 − 180 and that they essentially anticipate all the PBs in the network, as shown in Fig. 1.3 B. In more details, one observes that besides firing in a precise order the three critical neurons always fire within a narrow temporal window one after the other before a PB as shown in Fig. 1.3 A, (in particular see the blue cross correlation functions). Namely, *c*_1_ fires after *c*_0_ with an average time delay *ΔT*_c_1_,c_0__ ≃ 13.4 ms, and then *c*_2_ after *c*_1_ within the average interval *ΔT*_*c_2_,c_2_*_ ≃ 7.0 ms. Furthermore, the analysis of the activity of these three neurons during the inter-bursts reveal that they do not show anymore an univoque firing order or an unique activation time window (see in Fig. 1.3 A the red cross correlations functions).

**Fig. 1.2.**
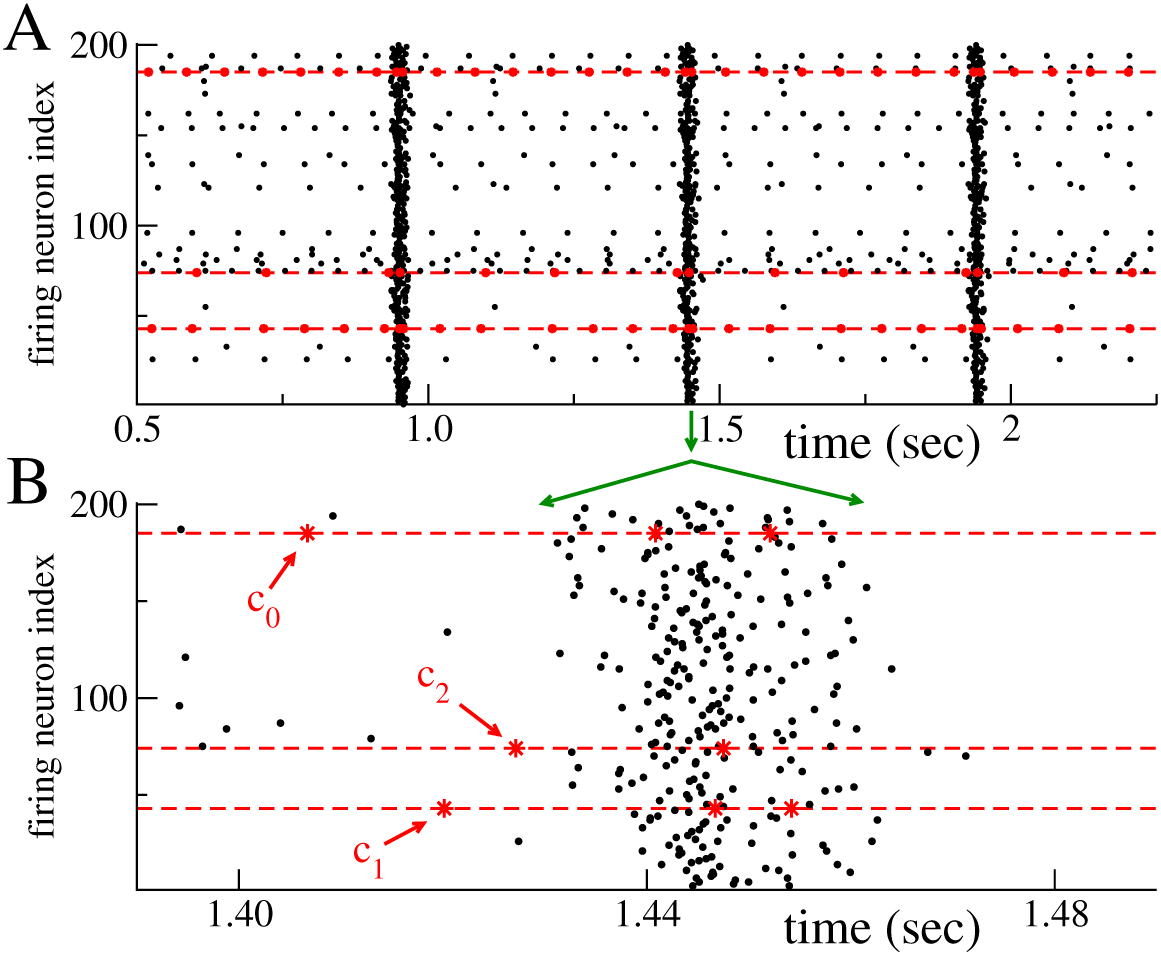
(A) Raster plot of the network activity: every dot signals a firing event. The (red) dashed lines and (red) stars refer to the critical neurons. (B) Close up of a population burst: PBs are anticipated by the ordered firing sequence of the critical neurons *c*_0_ → *c*_1_ → *c*_2_. In the raster plots, at variance with all the other figures, the neuronal labels are not ordered accordingly the ascending firing rates.

**Fig. 1.3.**
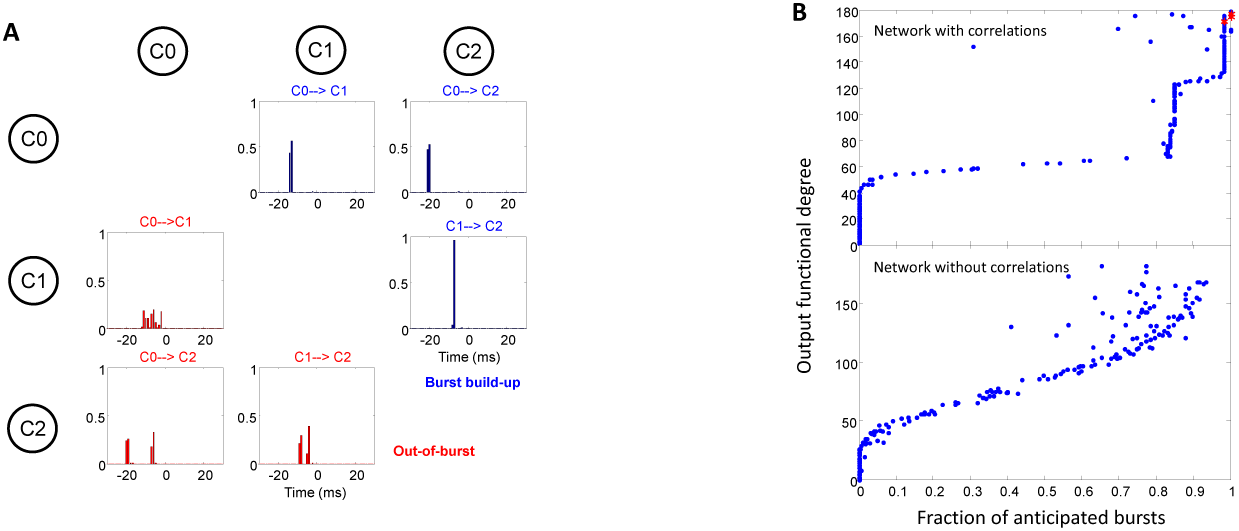
(A) Cross correlation functions *C*(*τ*) between the spike trains of two critical neurons for the network with correlations. The plots show all the possible pair combinations of the critical neurons: blue (red) histograms refer to the analysis performed during the population burst build up (during periods out of the bursting activity). The order of activation of each pair is reported on the top of the corresponding plot, whenever the cross-correlation has a significant maximum at some finite time *τ*_*max*_. (B) Output functional degree as a function of the fraction of the anticipated population bursts for all the neurons in the network: each dot denotes a different neuron. The data in the top (bottom) panel refer to the network with correlations (without correlations). The (red) stars on the top right in the top panel signal the critical neurons *c*_0_, *c*_1_, *c*_2_.

An extensive investigation of the critical neurons subjected to stimulations with currents in the range *I*^stim^ ∈ [14.5 : 16.0] mV reveals that PBs can be observed only if the neurons *c*_1_ and *c*_2_ had excitabilities within a narrow range (of amplitude ≃ 0.2 mV) centered around the threshold value. While, the stimulation of neuron *c*_0_ reveals that the network is always active for *I*^stim^ > *V*_th_, apart the occurence of an absolute minimum in the PB activity (an *anti-resonance*) at *I*^stim^ = 15.32 mV and a relative minimum at *I*^stim^ = 15.45 mV (see Fig. 1.4).

**Fig. 1.4.**
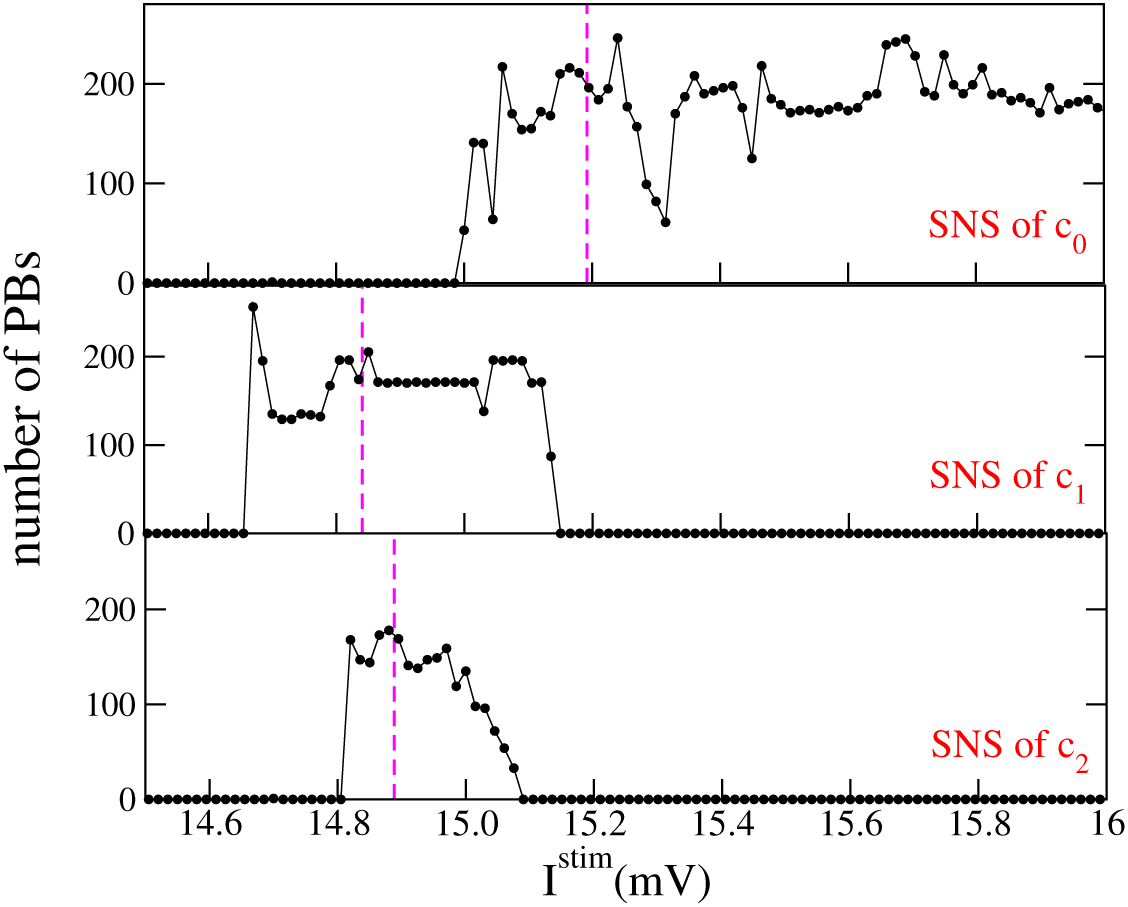
Number of PBs emitted in a time window *Δt* = 84 s during SNS experiments of the critical neurons *c*_0_, *c*_1_, *c*_2_ versus the stimulation current *I* ^stim^. The vertical dashed lines signal the value of the intrinsic excitability in control condition.

As shown in Fig.1.4, whenever *c*_1_ and *c*_2_ are stimulated with currents 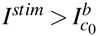 = 15.19 mV the bursting activity stops. This behaviour indicates that neuron *c*_0_ is the *leader* of the clique and the other ones are simply *followers* in the construction of the PB, they cannot fire more rapidly then neuron *c*_0_ or the PBs ceases. An analysis of the structural connectivity reveales that neuron *c*_0_ projected an efferent synapse on *c*_1_, which projected on *c*_2_.

The critical neurons, as already mentioned, are not hubs in a structural sense, since they have a very low connectivity *K^T^*, however they are indeed functional hubs, as shown from the previous analysis. Therefore, we can affirm that neurons *c*_0_, *c*_1_ and *c*_2_ form a functional clique, whose sequential activation, within the previously reported time windows, triggers the population burst onset.

The functional relevance of the neurons for the network dynamics is even more evident in other examples of functional cliques, reported in [13], where some of the supra-threshold neurons were even not structurally connected among each other.

## 1.4 Discussion

We have shown how, in a simple neural circuit displaying collective oscillations, the inclusion of developmentally inspired correlations among the single neuron excitabilities and connectivities can lead to the emergence of a *functional clique*. This is a small group of functionally connected neurons whose activation can be crucial to promote/arrest the collective firing activity in neural networks, irrespective of the underlying network topology. The clique is composed of a leader neuron, which can start the activation sequence at any moment, but to ignite the population burst the other two neurons, the followers, should fire in a precise order and with quite defined delays.

These results besides being of extreme interest for the neuroscience and dynamical system communities, pave the way for a new approach to the control of the dynamics of neural circuits. Coherence or decoherence in the network activity can be induced by proper stimulation protocols of few peculiar neurons, a subject of extreme interest in particular for the treatment of Parkinson disease with adaptive deep brain stimulation [34].

Future developments towards a more realistic neural circuit would require the extension of the model to include inhibitory neurons, as well as facilitation mechanisms at the level of synaptic transmission, and of network topology arising from anatomical data.

## Acknowledgements

We thank Y. Ben-Ari, D. Angulo-Garcia, R. Cossart, A. Malvache, L. Módol-Vidal, for extremely useful interactions. This article is part of the research activity of the Advanced Study Group 2016 *From Microscopic to Collective Dynamics in Neural Circuits* performed at Max Planck Institute for the Physics of Complex Systems in Dresden (Germany).

## References

1. Friston KJ (1994) Functional and effective connectivity in neuroimaging: a synthesis. Human brain mapping 2: 56–78.

2. Bullmore E, Sporns O (2009) Complex brain networks: graph theoretical analysis of structural and functional systems. Nature Reviews Neuroscience 10: 186–198.

3. Feldt S, Bonifazi P, Cossart R (2011) Dissecting functional connectivity of neuronal micro-circuits: experimental and theoretical insights. Trends in Neurosciences 34: 225–236.

4. Bullmore E, Sporns O (2012) The economy of brain network organization. Nature Reviews Neuroscience 13: 336–349.

5. Lee WCA, Reid RC (2011) Specificity and randomness: structure–function relationships in neural circuits. Current Opinion in Neurobiology 21: 801–807.

6. Boccaletti S., Latora, V., Moreno, Y., Chavez, M., and Hwang, D. U. (2006). Complex networks: Structure and dynamics. Physics reports, 424(4), 175–308.

7. Olmi S., Livi, R., Politi, A., and Torcini, A. (2010). Collective oscillations in disordered neural networks. Physical Review E, 81(4), 046119.

8. Luccioli, S., Olmi, S., Politi, A, and Torcini, A. (2012). Collective dynamics in sparse networks. Physical review letters, 109(13), 138103.

9. Gaiteri C, Rubin JE (2011) The interaction of intrinsic dynamics and network topology in determining network burst synchrony. Frontiers in Computational Neuroscience 5: 10.

10. Miller, J. E. K., Ayzenshtat, I., Carrillo-Reid, L., and Yuste, R. (2014). Visual stimuli recruit intrinsically generated cortical ensembles. Proceedings of the National Academy of Sciences, 111(38), E4053–E4061.

11. Seabrook, T. A., and Huberman, A. D. (2015). Cortical Cliques: A Few Plastic Neurons Get All the Action. Neuron, 86(5), 1113–1116.

12. Barnes, S. J., Sammons, R. P., Jacobsen, R. I., Mackie, J., Keller, G. B., and Keck, T. (2015). Subnetwork-specific homeostatic plasticity in mouse visual cortex in vivo. Neuron, 86(5), 1290–1303.

13. Luccioli, S., Ben-Jacob, E., Barzilai, A., Bonifazi, P., and Torcini, A. (2014). Clique of functional hubs orchestrates population bursts in developmentally regulated neural networks. PLoS Comput Biol, 10(9), e1003823.

14. Tsodyks M., Kenet, T., Grinvald, A., and Arieli, A. (1999). Linking spontaneous activity of single cortical neurons and the underlying functional architecture. Science, 286(5446), 1943–1946.

15. Brecht M, Schneider M, Sakmann B, Margrie TW (2004) Whisker movements evoked by stimulation of single pyramidal cells in rat motor cortex. Nature 427: 704–710.

16. Houweling AR, Brecht M (2007) Behavioural report of single neuron stimulation in somatosensory cortex. Nature 451: 65–68.

17. Cheng-yu TL, Poo Mm, Dan Y (2009) Burst spiking of a single cortical neuron modifies global brain state. Science 324: 643–646.

18. Wolfe J, Houweling AR, Brecht M (2010) Sparse and powerful cortical spikes. Current Opinion in Neurobiology 20: 306 - 312.

19. London M, Roth A, Beeren L, Häusser M, Latham PE (2010) Sensitivity to perturbations in vivo implies high noise and suggests rate coding in cortex. Nature 466: 123–127.

20. Kwan AC, Dan Y (2012) Dissection of cortical microcircuits by single-neuron stimulation in vivo. Current Biology 22: 1459–1467

21. Bonifazi P, Goldin M, Picardo MA, Jorquera I, Cattani A, et al. (2009) Gabaergic hub neurons orchestrate synchrony in developing hippocampal networks. Science 326: 1419–1424.

22. Allène C, Cattani A, Ackman JB, Bonifazi P, Aniksztejn L, et al. (2008) Sequential generation of two distinct synapse-driven network patterns in developing neocortex. The Journal of Neuroscience 28: 12851–12863.

23. Tsodyks M, Uziel A, Markram H (2000) Synchrony generation in recurrent networks with frequency-dependent synapses. The Journal of Neuroscience 20: 50RC+.

24. Tsodyks MV, Markram H (1997) The neural code between neocortical pyramidal neurons depends on neurotransmitter release probability. Proceedings of the National Academy of Sciences 94: 719–723.

25. Turrigiano GG, Leslie KR, Desai NS, Rutherford LC, Nelson SB (1998) Activity-dependent scaling of quantal amplitude in neocortical neurons. Nature 391: 892–896.

26. Turrigiano GG (2008) The self-tuning neuron: synaptic scaling of excitatory synapses. Cell 135: 422–435.

27. Ge S, Goh EL, Sailor KA, Kitabatake Y, Ming Gl, et al. (2005) Gaba regulates synaptic integration of newly generated neurons in the adult brain. Nature 439: 589–593.

28. Doetsch F, Hen R (2005) Young and excitable: the function of new neurons in the adult mammalian brain. Current Opinion in Neurobiology 15: 121–128.

29. Marissal T, Bonifazi P, Picardo MA, Nardou R, Petit LF, et al. (2012) Pioneer glutamatergic cells develop into a morpho-functionally distinct population in the juvenile ca3 hippocampus. Nature Communications 3: 1316.

30. Picardo MA, Guigue P, Bonifazi P, Batista-Brito R, Allene C, et al. (2011) Pioneer gaba cells comprise a subpopulation of hub neurons in the developing hippocampus. Neuron 71: 695–709.

31. Karayannis T, García NVDM, Fishell GJ (2012) Functional adaptation of cortical interneurons to attenuated activity is subtype-specific. Frontiers in Neural Circuits 6: 66.

32. Bonifazi P, Difato F, Massobrio P, Breschi GL, Pasquale V, et al. (2013) In vitro large-scale experimental and theoretical studies for the realization of bi-directional brain-prostheses. Frontiers in Neural Circuits 7: 40.

33. Ben-Ari Y (2002) Excitatory actions of gaba during development: the nature of the nurture. Nature Reviews Neuroscience 3: 728–739.

34. Little S., et al. (2013). Adaptive deep brain stimulation in advanced Parkinson disease. Annals of neurology, 74: 449–457.

